# Modeling invasion patterns in the glioblastoma battlefield

**DOI:** 10.1101/2020.06.17.156497

**Authors:** Martina Conte, Sergio Casas-Tintò, Juan Soler

## Abstract

Glioblastoma is the most aggressive tumor of the central nervous system, due to its great infiltration capacity. Understanding the mechanisms that regulate the Glioblastoma invasion front is a major challenge with preeminent potential clinical relevance. In the infiltration front, the key features of its dynamics relate to biochemical and biomechanical aspects, which result in extended cellular protrusions, known as tumor microtubes. The coordination of metalloproteinase expression, extracellular matrix degradation, and integrin activity emerges as leading mechanism that facilitates Glioblastoma expansion and infiltration in uncontaminated brain regions. We propose a novel multidisciplinary approach, based on in vivo experiments in *Drosophila* and mathematical models, for the proteins dynamics at the front of Glioblastoma, with a predictive value of the tumor progression.

## Introduction

Glioblastoma (GB) is the most common, aggressive, and lethal tumor of the central nervous system. It has glial origin and it is characterized by rapid cell proliferation, great infiltration capacity, and neurological impairment [1]. GB is composed of a heterogeneous genetic landscape of tumor cells [2], which reduces the efficiency of clinical treatments. Current treatments for GB include surgical resection of the solid tumor, radiation therapy, and chemotherapy with Temozolomide. However, GB is resistant to treatment and recurrence is the main cause of mortality [3, 4]; the median survival is below16 months and the incidence is 3/100.000 per year.

GB infiltration in the human brain is a complex phenomenon, influenced by different tumor properties and the tumor microenvironment, including brain-resident cells, blood brain barrier, and immune system [5]. In particular, GB develops fronts of invasion towards uncontaminated areas of the brain, and GB cells modify their cytoskeleton components [6] to extend protrusions, known as Tumor Microtubes (TMs) [7]. TMs mediate GB progression, and they interact with synapses of neighboring neurons [8, 9]. But, how do these fronts occur? Which are the mechanical and chemical processes characterizing these front areas of GB progression? Which are the main agents involved? What is the role of TMs in the tumor front development? Is there any heterogeneity in the spatial distribution of the tumor activity across the tumor domain?

These are some of the questions that we address throughout the paper, providing insights into the role of integrins, metalloproteases, and TMs in GB growth and spread. Our results are supported by experimental measurements in a *Drosophila* model of human GB, in continuous feedback with an evolutionary mathematical model that predicts behaviors and interactions of these biological agents.

### GB in *Drosophila*

The most frequent genetic lesions in GB include the constitutive activation of phosphatidylinositol 3-kinase (PI3K) and Epidermal Growth Factor Receptor (EGFR) pathways, which drive cellular proliferation and tumor malignancy [10]. *Drosophila melanogaster* has emerged as one of the most reliable animal models for GB; it is based on equivalent genetic mutations in EGFR and PI3K pathways found in patients [11]. Glial cells respond to this oncogenic transformation and reproduce all main features of the disease, including glial expansion and invasion, but in a shorter period of time. This *Drosophila* model has been used for drug and genetic screenings and the results have been validated in human GB cells (REFs) [12, 13, 14].

### Agents involved in the battlefield

GB growth and migration are driven by specific signaling pathways as well as interactions between the tumor and its extracellular microenvironment. In our study, we include the cell membrane protrusions (TMs) as driving factors of tumor progression, as well as cell response to signaling gradients and the interplay with the Extra Cellular Matrix (ECM). A schematic diagram of the described dynamics is given in Figure 1.

**Figure 1:**
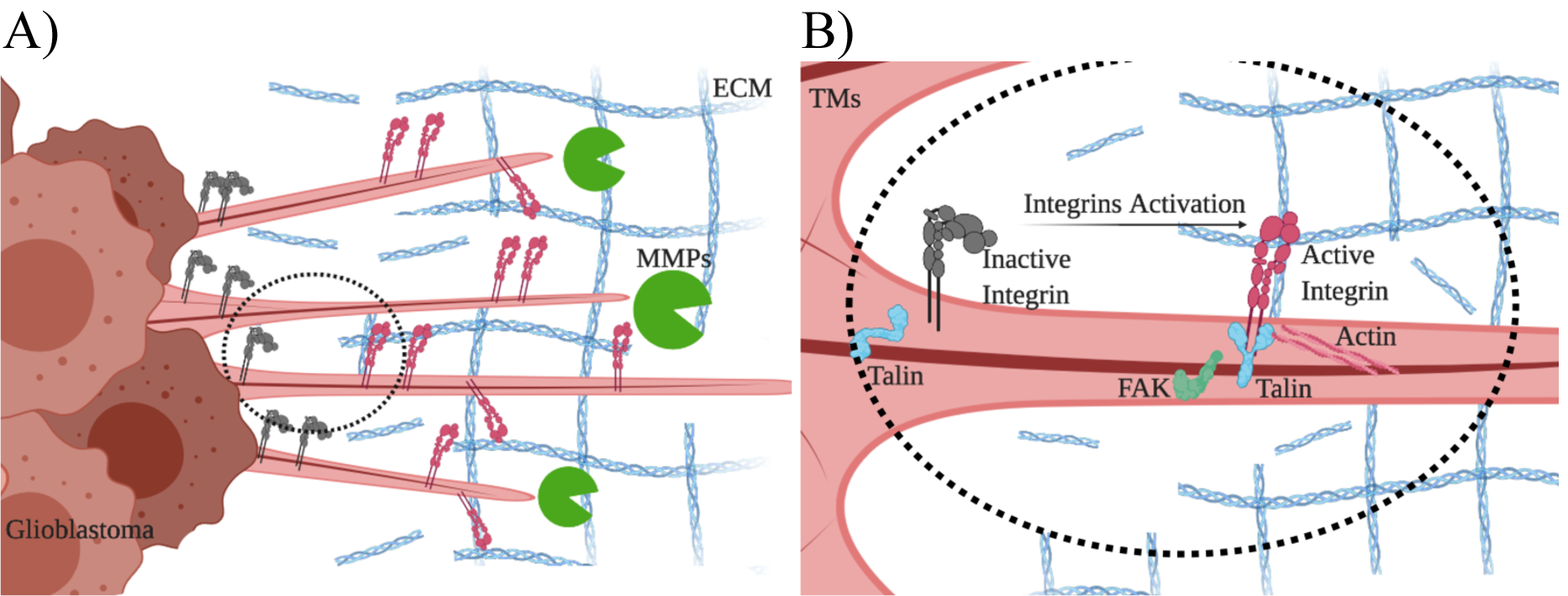
Diagram of GB and proteins dynamics. Diagram of the interactions between the proteins involved in GB dynamics and progression, which give rise to the mathematical model. A): GB cells produce and release in the extracellular space Matrix Metalloproteinases (MMPs) that proteolyze Extra Cellular Matrix (ECM) components. B): Magnification of TMs. Integrins are activated in the GB Tumor Microtubes (TMs) upon interaction with ECM proteins. Active Integrins, interacting with Actin filaments and Talin adaptor protein, activate FAK protein to promote cytoplasm dynamics.

The dynamics of tumor cell membrane, including cell protrusions, is fundamental in several processes, among others cell movements, cell transport, and exposure to molecular interactions with substrates during GB progression. Generally, cell protrusions are highly dynamic extensions of the plasma membrane that are involved in cell migration and invasion through the ECM. Different types of protrusions have been identified to contribute to cell spreading, depending on specific contexts, cell types, and microenvironment. In particular, filopodia are thin, finger-like and highly dynamic membrane protrusions that have a significant role in mediating intercellular communication and in modulating cell adhesion. They appear to be required for haptotaxis and chemotaxis [15], i.e., for the cell response to the gradient of insoluble (haptotaxis) and soluble (chemotaxis) components of the tumor microenvironment. Cytonemes are one type of filopodia, first observed in the *Drosophila* wing imaginal disc [16] (see also [17, 18, 19, 20, 21, 22, 23]); also known as TMs in the context of GB, they have an important role in tumor development [12]. Specifically, we focus on TM involvement in the GB front progression and on their relationship with integrins and proteases.

Tumor propagation is characterized by a sharp invasion front located in the front area of GB progression, where TMs are mostly concentrated. The agents involved in GB invasion, such as integrins, proteases or the tumor cells themselves, are neither scattered nor randomly moving, but rather there is a self-organization that determines invasion patterns around the tumor front. This crucial aspect for the progression of GB can be observed in Figure 2, where proteases and integrins co-localize with the TM region at the GB front.

**Figure 2:**
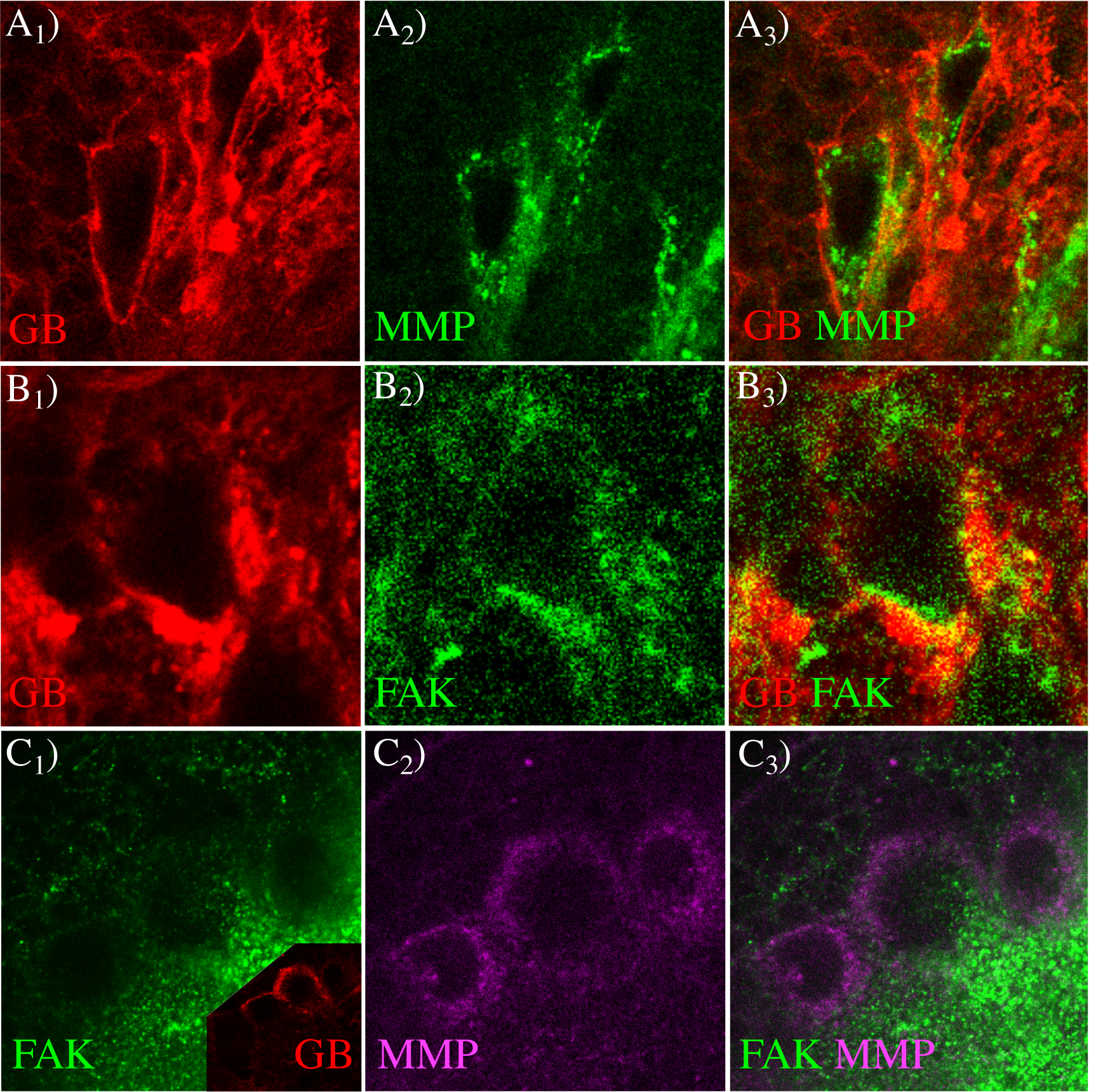
MMP1 and FAK proteins in GB. Fluorescent confocal images of *Drosophila* 3rd instar larvae brain with GB. *A*) Glial membrane is shown in red (*A*_1_), and MMP1 protein in green (*A*_2_). MMP1 accumulation in GB front is observed in merged images (*A*_3_). *B*) Glial membrane (*B*_1_, red) and FAK distribution (*B*_2_, green) signals are shown through the brain. FAK accumulation in the GB front is detailed in merged images (*B*_3_). *C*) Co-staining of FAK (*C*_1_, green) and MMP1 (*C*_2_, magenta), and merged images (*C*_3_). FAK and MMP1 signals accumulate in the same region of GB front (inset in *C*_1_ red).

The prediction of these expansion patterns is a need to envisage possible outcomes in patients and for the development of potential therapies. Therefore, any mathematical model that attempts to predict GB dynamics and reproduce the formation of these evolutionary patterns must face these challenges.

Integrins are transmembrane glycoproteins known to mediate dynamic interactions between the ECM and the actin cytoskeleton during cell motility. The binding specificity is determined by the integrin extracellular domain, that allows the recognition of different matrix ligands (fibronectin, collagen, laminin) and other cell surface receptors [24]. In *Drosophila*, the gene *rhea* encodes Talin, a large adaptor protein required for integrin function. Talin links clusters of integrins, bound to ECM, and the cytoskeleton ant; therefore, it is essential for the formation of focal adhesion-like clusters of integrins [25]. Integrin interactions with ECM components support cell adhesion, spread and migration [26], affecting also cell growth and division through the additional interaction with extracellular proteins and enzymes that control cell cycle [27, 28]. Integrins are generally present in the cellular membrane, but mostly at the tumor front, involving directly their receptors in the signaling leading the migration process.

Proteases are enzymes that catalyze the breakdown of proteins into smaller polypeptides or single amino acids. Some type of proteases are localized around the cell membrane and play a key role in promoting tumor invasion and tissue remodeling. They induce proteolysis of ECM components [29] and maintain a microenvironment that facilitates tumor cell survival. Protease profiling studies have indicated that the expression of serine proteases, cysteine proteases and matrix metalloproteases (MMPs), the most prominent family of proteinases associated with tumorigenesis, increase in high-grade astrocytoma compared with normal brain. In particular, increased MMP levels and tumor invasiveness in human GBs show a strong correlation [30, 31, 12]. Several studies reported their localization in TM membrane [32, 33], especially for MMP-2 and MT1-MMP.

Our interest arises from the analysis of the role of TMs in tumor progression and of the effect of different drugs on their structures and dynamics [34, 35]. The inhibition of TMs leads to a decrease in cell migration and proliferation, with obvious consequences on GB treatment [34, 35, 36]. The process of growth and retraction of TMs plays a fundamental role for cell-to-cell communication routing [17, 18, 20, 21, 22, 23], as well as a transient binding platform for essential proteins that regulate tumor dynamics [37]. Among these proteins, the presence of microtubule plus end-binding protein EB1 correlates with GB progression and poor survival [36, 38]. This has led to a great interest in the development of drugs aimed at affecting end-binding proteins expression and, consequently, TM dynamics [34]. Moreover, several experiments have reported the influence of proteases on GB development and the role of several integrin receptors in its progression (see [39, 40] and Supplementary Texts S1 and S3 for further details).

### Predictions and modeling

The ability of mathematical models to predict and guide biological experiments has gathered great attention. In fact, the number of possible factors involved in GB progression is large and the relevance of each of them is still unknown. Thus, exploiting the advances in microscopy and protein signaling, there is a need to integrate the large amount of provided data into mathematical models.

Different mathematical models describe the evolution of tumors by linear (Fick’s law) or nonlinear diffusion (Darcy’s law); some approaches couple dynamics for tumor cells and environmental agents involved in the tumor evolution, some others consider them independently (see [41, 42] and references therein). However, the arising of sharp profiles is not compatible with a movement induced by linear diffusive dynamics, although some mathematical models explored this possibility as a first approach.

We focus on the dynamics of the tumor propagation front and on the emergence of the coordination between self-organized collective processes. This coordination is strongly non-linear and the different agent dynamics adapt to each other. The propagation front arises and locates in an area that occupies about 5-7 cell diameters ahead of the tumor main body, where we report an enhancement of tumor activity and integrin binding (see Figure 2).

We present here a novel mathematical model that covers specific evolutionary aspects of tumor progression. From the modeling side, the novelties of our approach include the description of the dynamics of the tumor front, and the link of cell membrane movements with the dynamics of some proteins, their concentration and location in the TM region. In particular, tumor density is governed by haptotactic and chemotactic processes, induced by MMP and active integrins, as well as by a flux saturated mechanism (see [43, 44, 17] and references therein). The latter allows the incorporation of biological features related to tumor dynamics (i.e., the viscosity of the medium and the speed of propagation) and the definition of sharp invasion profiles [43]. Therefore, this model acquires an excellent predictive capability, both quantitatively and qualitatively, since advanced mathematical concepts are confluent with precise biological experiments and, furthermore, the predictions of our model have partly guided the experimental research, which proves its consistency.

## Results

### Localizing the front of GB

We focus on the dynamics of the active front in GB. In particular, we analyze the distributions of MMP1, integrins and focal adhesions and their localization in relation with the tumor membrane density. We hypothesize a spatial het-erogenous activity of tumor cells, in terms of proteolysis and binding receptor activation. This spatial heterogeneity might determine the accumulation of MMP1 and the activation of focal adhesions at the tumor front.

First, we show the presence of MMP1 protein at the active front of GB tissue. We dissected *Drosophila* brain samples with a genetically induced GB (*repo*>*PI3K; EGFR*). To delimitate the front of the tumor, we induced the co-expression of a membrane bound (myristoylated form) version of the red fluorescent protein (*UAS-myrRFP*) and, accordingly, all GB cell membranes were marked in red. MMP1 protein was detected by a specific monoclonal anti-MMP1 antibody (Figure 2-*A*_2_ green). The confocal microscopy images show that MMP1 signal is heterogeneous through the brain, and stronger in specific regions of GB front (Figure 2-A red). Then, we visualized focal adhesions with a specific monoclonal antibody anti-FAK (Figure 2-*B*_2_ green). The confocal images show that FAK staining concentrates in the GB front mostly. Finally, we co-stained for FAK (Figure 2-*C*_1_ green) and MMP1 (Figure 2-*C*_2_ magenta) proteins, and visualized the active front of GB cells (Figure 2-*C*_1_ red inset). The confocal fluorescent images show that MMP1 and active focal adhesion (FAK) signals coincide at the tumor front. These results are compatible with our hypothesis that MMP1 and FAK are characteristics features of the leading edge of migrating cells, with a central role in the process of GB expansion.

Therefore, we model the total flux of tumor cells (denoted by 𝒥_*N*_ (*N, P, A*)) by a combination of three main factors. Firstly, the dynamics that 𝒥_*N*_ exerts on cell movement include a saturated flow [44] (see Supplementary Text S1 for a detailed description). It allows to define the movement of a sharp (non-diffusive) profile [43] and to incorporate the experimental data about the propagation front velocity and the porosity of the medium. Then, 𝒥_*N*_ collects information about the cell response to the gradient of soluble and insoluble components of the tumor microenvironment. In particular, we consider gradients of MMP1 (*P* (*t, x*)) for the chemotactic process, and integrins (*A*(*t, x*)) for the haptotactic process, nonlinearly coupled with the concentration of GB cells. Mathematically, the dynamics of the tumor cells density (*N* (*t, x*)) is modeled as:

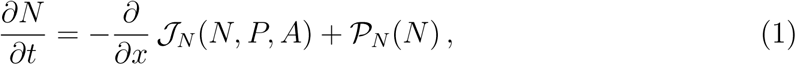

where, in addition to the total flux 𝒥_*N*_ (*N, P, A*), we use the term 𝒫_*N*_ (*N*) to describe tumor proliferation.

Using this modeling approach, we demonstrate not only the role of MMP1 and integrins in GB motility, but also their co-localization and spatial distributions with respect to the location of the tumor front, a region characterized by a higher tumor membrane density compared to the bulk tumor (region of higher tumor density).

### MMP1s distribution

MMP1s facilitate the tumor invasion process by degrading the extracellular matrix. These proteins are produced by GB cells and released in the extracellular space, mostly in the area around the tumor invasion front. Here, TMs are present in large numbers, thus, enhancing the proteolytic activity. Therefore, we propose the localization of MMP1s on TMs and a mathematical modeling of MMP1 evolution in relation to the tumor front.

Experimentally, to analyze the protein distribution in the tumor, we quantified the MMP1 signal in the inner GB mass and at the GB border. We visualized *Drosophila* brains with GB (Figure 3-*A*_1_ and 3-*B*_1_) and immunostained with anti-MMP1 (Figure 3-*A*_2_ and 3-*B*_2_). The results indicate that, at the front region of GB (Figure 3-*A*_1_) where the membrane density (red in Figure 3-*A*_3_) is higher, MMP1s (green in Figure 3-*A*_3_) accumulate, showing a peak of concentration in the region corresponding to the TMs. Interestingly, this MMP1 maximum appears slightly shifted in the direction of GB migration with respect to the peak of GB membrane density. We analyzed the inner GB mass by taking a wider region of GB brains to measure MMP1 and GB density. The confocal images show homogeneous lower levels (basal levels) of GB membrane and MMP1 protein in the inner region (Figure 3-*B*_1_-*B*_3_) compared to the front (Figure 3-*A*_1_- *A*_3_). That is, MMP1 protein accumulation occurs specifically in the front region of GB, correlating with the peaks of GB cell membrane density in the TM region.

**Figure 3:**
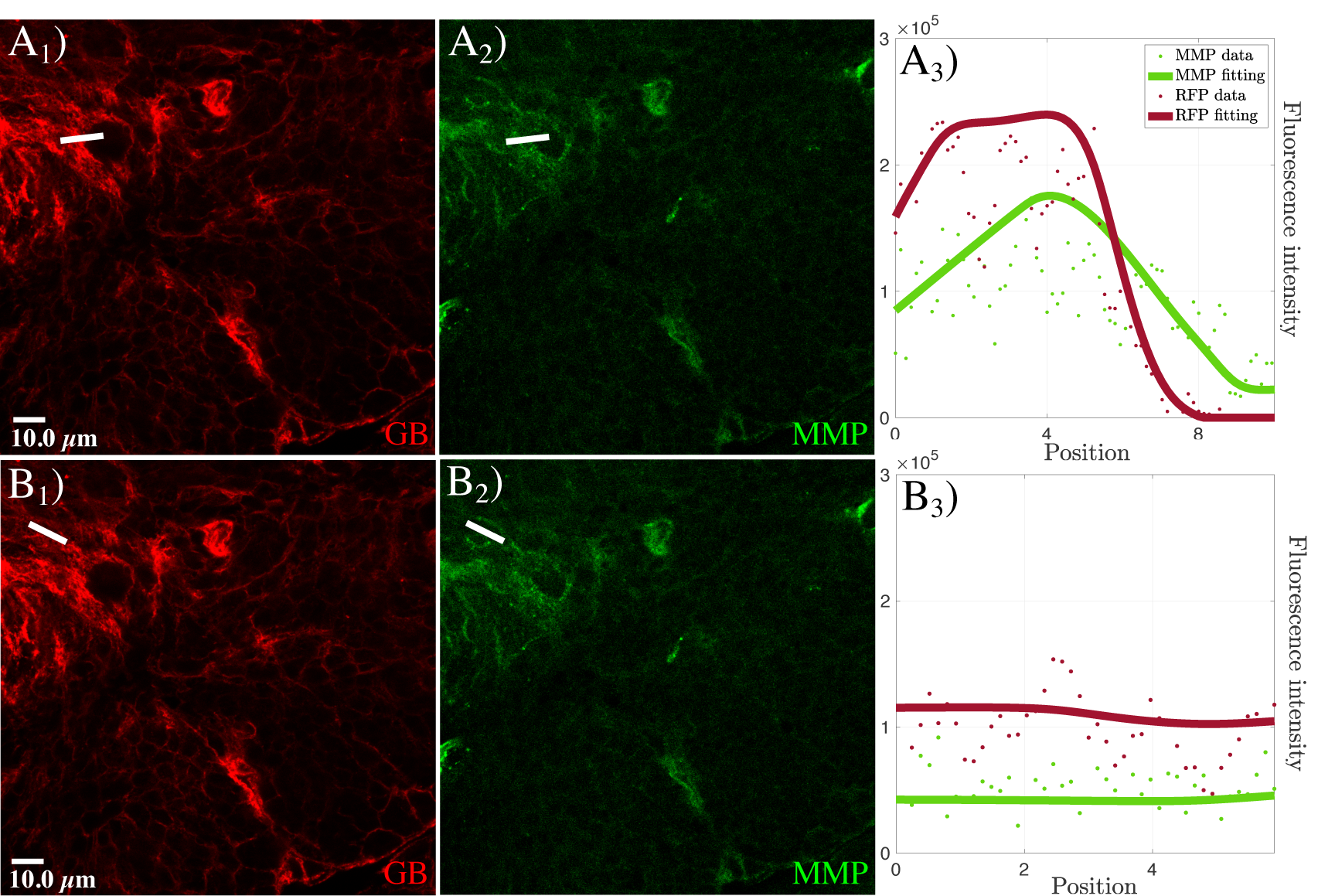
MMP1 accumulation in GB front and inner GB mass. Fluorescent confocal images of *Drosophila* 3rd instar larvae brain with GB marked in red (white line in *A*_1_ is front and white line in *B*_1_ is inner GB mass), and stained with anti-MMP1 in green (*A*_2_ and *B*_2_). In *A*_3_ and *B*_3_, quantification and graphical representation of the fluorescent intensity for GB and MMP1 signals along the white lines in *A*_1_, *A*_2_, *B*_1_ and *B*_2_. Dots represent the data and lines represent the fitting.

We model the dynamics and the distribution of MMP1s (*P* (*t, x*)) by taking into account three main phenomena, represented in the following equation:

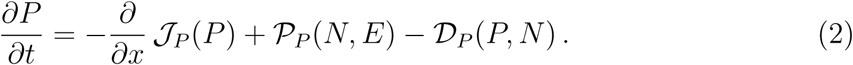

MMP1 production (𝒫_*P*_) by tumor cells is localized along the TM region, where the neoplastic tissue is in contact with the healthy tissue, and it is directly dependent on the heterogeneous tumor proteolytic activity. After MMP1 release in the extracellular space, its flux (𝒥_*P*_) is limited in the areas surrounding the tumor mass, generating a sharp front ahead of the tumor. As the tumor front advances, the remaining MMP1 is degraded (𝒟_*P*_) and maintains basal levels in the inner tumor region (further details are given in the Supplementary Text S1).

MMP1 evolution process determines a dynamical degradation of the ECM (*E*(*t, x*)), described as:

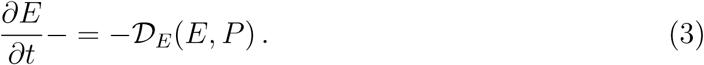

### Focal adhesions and integrin dynamics

Integrins are transmembrane receptors that allow cells to bind ECM ligand facilitating tumor cell movement. Upon activation, integrins mediate the organization of the cytoskeleton, cell cycle, and cell migration. In GB cells, the activation process occurs predominantly in the TM region and active integrins result homogeneously distributed throughout TMs. Moreover, the conversion into the inactive (no bound) state occurs in the proximal region of the TMs with respect to the main GB mass, as shown by our experimental and mathematical results.

To determine the molecular changes at the GB front in relation with the activity of integrins, we immunostained GB brain samples with Talin, a mediator of integrin adhesivity [45], and with Focal Adhesion Kinase (FAK), a cytoplasmic Tyrosine kinase involved in signaling and cytoskeleton dynamics associated to integrin activity [46, 47]. We analyzed confocal microscopy images of GB front regions, and compared fronts with low or high GB membrane signal.

The quantification of the signals for anti-Talin and anti-FAK (Figure 4-*A*_1_-*B*_4_) shows that Talin (black in Figure 4-*A*_4_ and 4-*B*_4_) is reduced in the TM region, i.e., where the GB membrane signal is high, while FAK (magenta in Figure 4-*A*_4_ and 4-*B*_4_) is increased, thus the pattern of Talin and FAK expression is inverted at the front. Additionally, both GB membrane and FAK signals correlate. The GB/FAK signal correlation is maintained in different fronts, irrespective of their low (Figure 4-*A*) or high (Figure 4-*B*) levels of GB membrane signal. These results suggest an equivalent correlation for the reduction of Talin and the increase of FAK signals at the GB front, consistent with the relation of integrin dynamics and cell motility.

**Figure 4:**
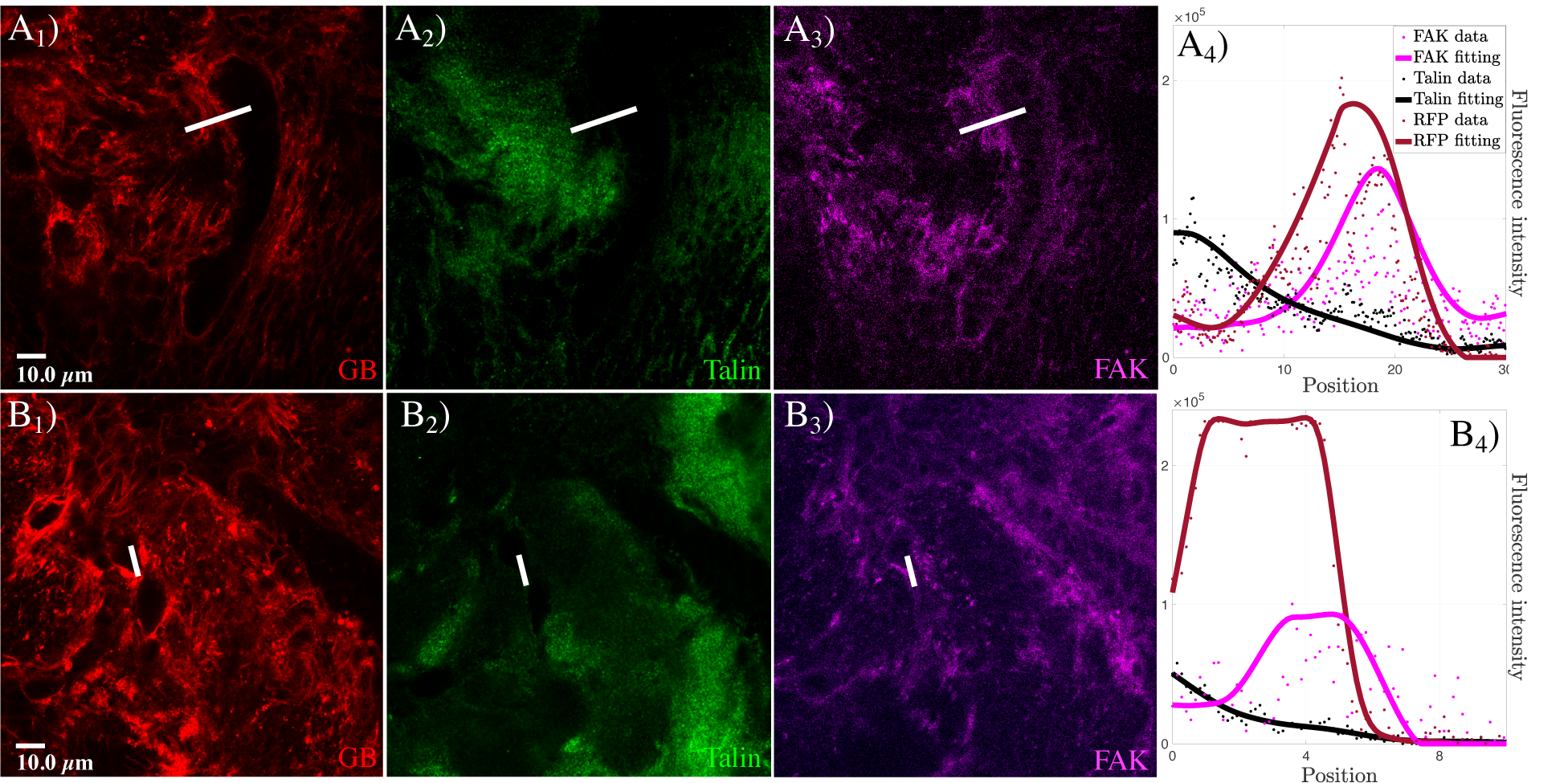
Talin and FAK dynamics in the transition between GB front and healthy tissue. Fluorescent confocal images of *Drosophila* 3rd instar larvae brain with GB marked in red (*A*_1_ and *B*_1_), and stained with anti-Talin (green) and anti-FAK (magenta) (*A*_2_, *A*_3_, *B*_2_ and *B*_3_). In *A*_4_ and *B*_4_, quantification of the fluorescent signals and graphical representation of the fluorescent intensity for GB, Talin and FAK signals along the white lines shown in the images *A*_1_-*A*_3_ and *B*_1_-*B*_3_. Dots represent the data and lines represent the fitting.

To confirm the functional contribution of integrins to GB progression and validate these suggestions, we used specific RNAi constructs to knockdown *myospheroid (mys)*, the *Drosophila* Integrin B subunit, or *rhea*, the *Drosophila* Talin, two key players for integrin function. The data show that GB cells require integrins to progress and expand (Supplementary Figure S1 A-C). Moreover, *mys* or *rhea* knockdown rescues the lethality caused by GB (Supplementary Figure S1-D). Finally, to confirm Talin and FAK inverse correlation, we compared inner GB mass and GB front areas. The data (Supplementary Figure S2) confirm that Talin and FAK maintain an inverse correlation, and they are indicators of the migratory status of the GB cells.

Mathematically, we split the integrin population into two subpopulations, referring to the active (*A*(*t, x*)) and the inactive state (*I*(*t, x*)) of the integrin receptors. Their dynamics take into account these activation, and consequent inactivation, processes:

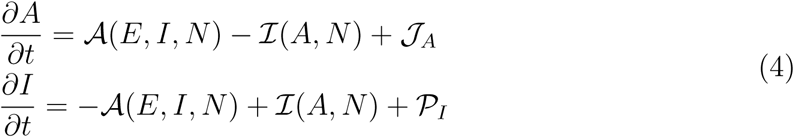

In particular, activation (𝒜(*E, I, N*)) is mediated by the presence of ECM, to which cells bind, and it depends on the heterogenous tumor activity on the TM region. Instead, the inactivation of integrins (ℐ(*A, N*)) happens once the tumor has crawled on the ECM and moved forward. Due to the movement of the tumor cells, a flux of active integrins (𝒥_*A*_) arises and influences the overall dynamics during crawling, with an estimated velocity depending on the tumor progression. Assuming that, initially, the inactive integrins along cell membrane have not reached the saturation value, a process of integrins production by exocytosis is also included in the model (*𝒫*_*I*_). In addition, considering the experimental results shown in the Supplementary Figure S2, a basal level of active integrins in the tumor main body is included, as well as a basal level of inactive integrins along the microtubules (for further details, see the Supplementary Text S1).

### Motility features of GB cells at the front

The results shown in Figure 2, regarding the features that characterize tumor front, together with the data on MMP1 and integrin dynamics, support our hypothesis on the processes that lead the motility of GB. Experimentally, we analyzed MMP1 concentration and focal adhesions dynamics to prove their role as cues for cell motility, invasiveness and migration [48, 49]. We co-stained GB brain samples with anti-MMP1 and anti-FAK (Figure 5) and quantified the intensity of the fluorescent signal. We compared the signal in low and high tumor density fronts (Figure 5-*A* and 5-*B* respectively).

**Figure 5:**
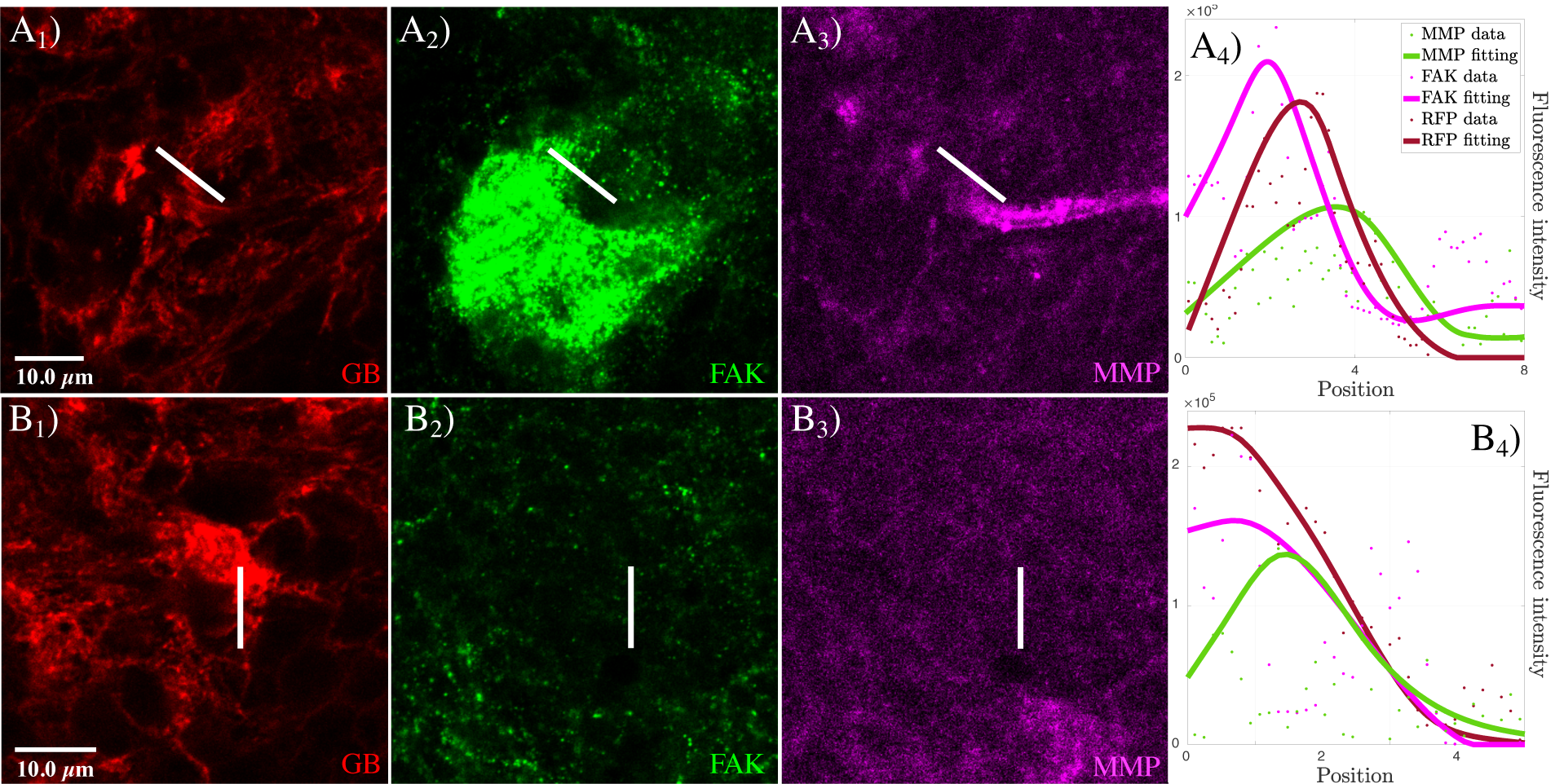
FAK and MMP dynamics in the transition between GB front and healthy tissue. Fluorescent confocal images of *Drosophila* 3rd instar larvae brain with GB marked in red (*A*_1_ and *B*_1_), and stained with anti-FAK (green) and anti-MMP1 (magenta) (*A*_2_, *A*_3_, *B*_2_ and *B*_3_). In *A*_4_ and *B*_4_, quantification of the fluorescent signals and graphical representation of the fluorescent intensity for GB, FAK and MMP1 signals along the white lines shown in the images *A*_1_-*A*_3_ and *B*_1_-*B*_3_. Dots represent the data and lines represent the fitting.

The results show that, at the GB front (Figure 5*A*_1_ and 5-*B*_1_), the maximum signal for FAK (Figure 5-*A*_2_ and 5-*B*_2_) occurs before the peak of MMP1 (Figure 5*A*_3_ and 5*B*_3_). This correlation occurs for both types of GB fronts, suggesting that this phenomenon is not a consequence of higher or lower GB membrane density. These results point towards a coordinated function between MMP1 activity and FAK dynamics, and suggest that, at the front, FAK acts closer to GB mass. By contrast, MMP1 plays its role further away from the front. Mechanistically, these results indicate that MMP1 activity in the proteolysis of the ECM is prior to the increase of integrin dynamics, it is associated to the presence of FAK, and, thus, together they contribute to the motility of GB cells.

### Modeling results

From a mathematical viewpoint, we numerically solve our model (the whole system (21) is provided in the Supplementary Text S1) with no flux boundary conditions. The results, represented in Figure 6, remarkably show how the model predicts that the region of greatest interest for the whole GB development process is the front, where TMs are located. The observed biological features correlate with the generation and evolution of this critical region of the tumor, the front.

**Figure 6:**
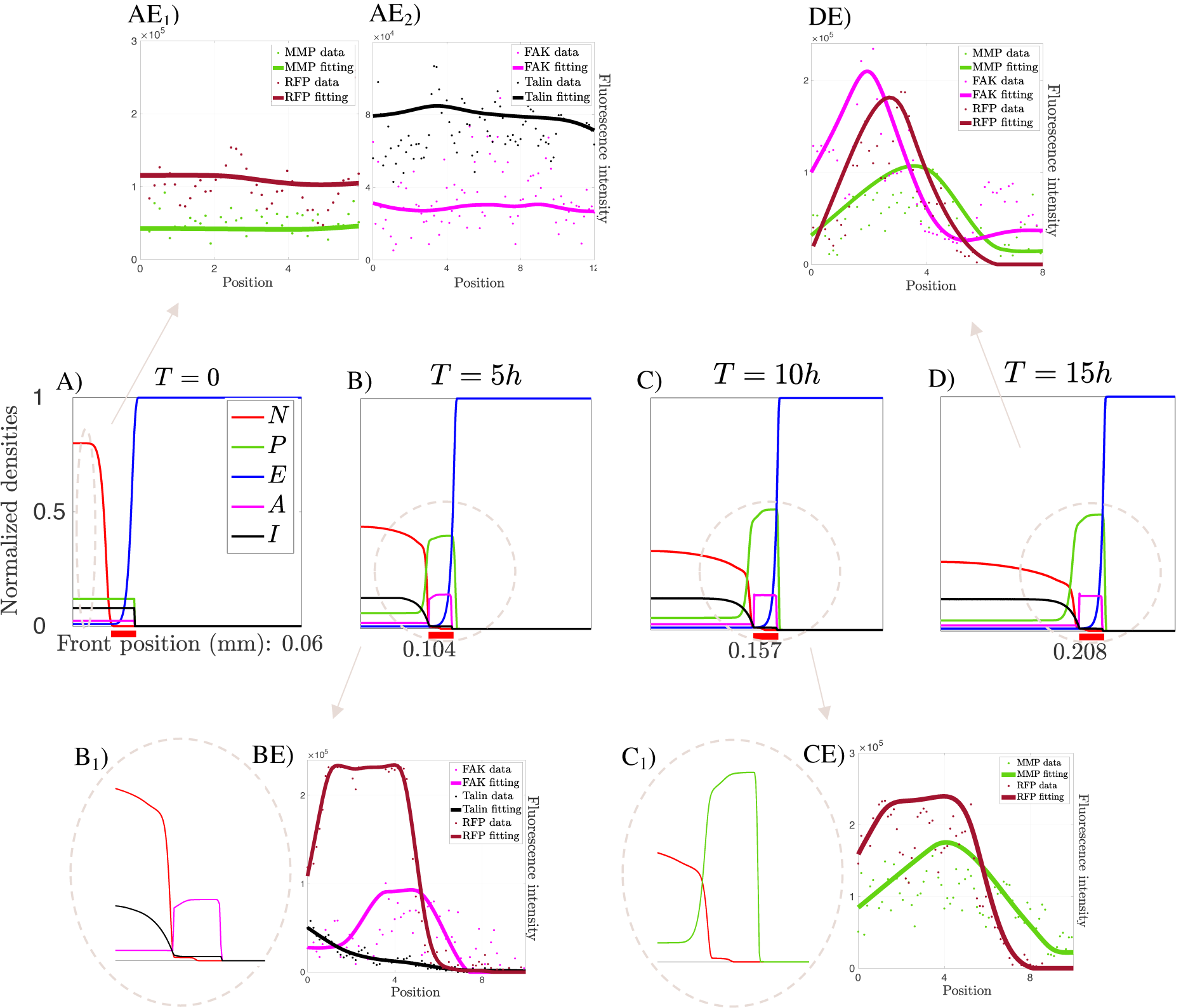
Modeling predictions. *A*), *B*), *C*) and *D*) are four snapshots of the numerical solution of our evolutionary model at the initial state, after 5, 10 and 15 hours, respectively. Images show the evolution of tumor density (*N*), ECM (*E*), active (*A*) and inactive (*I*) integrins and MMPs (*P*). The thick red line below the figures indicates the front area with the highest concentration of TMs. Images *AE*_*i*_), *BE*), *CE*) and *DE*) show the results of the analysis of the experimental data. In particular, peaks in the RFP distribution indicate areas with high tumor membrane density, i.e., the TM regions, while analogous peaks in the numerical simulations refer to high tumor density areas, i.e., the main tumor mass. The images *B*_1_) and *C*_1_) stand for a zoom of the indicated areas and they are correlated with the experimental data *BE* and *CE*, respectively. The parameters used in these simulations are listed in the Supplementary Table S1.

Figure 6-*A* shows the initial condition of the system, and it is accompanied by the quantifications of the basal levels of MMP1s with respect to the tumor density (Figure 6-*AE*_1_), and those of the active versus the inactive integrins (Figure 6 - *AE*_2_). In both cases, the measurements are done in areas inside the bulk tumor and the resulting data are incorporated into the model.

Figures 6-*B* and 6-*C* show the evolution at 5 and 10 hours, respectively, of the initial data given in Figure 6-*A*. The numerical simulations show that the model is capable of predicting, as specific invasion profiles, the evolution of the fronts for each of the agents involved in the migration process. Precisely, Figures 6-*B* and 6-*C* illustrate how the model collects the change between the active and inactive integrins at the beginning of the tumor front area, and how this corresponds to the experimental results (Figure 6- *BE*). A plateau-like profile arises for *A* in the TM region and, comparing Figure 6*B*_1_ with Figure 6-*BE*, these numerical results show a good agreement with the experimental data. Moreover, in these images the model predictions regarding the MMP1 front correlate with the results obtained experimentally, that are shown in Figure 6-*CE*. MMP1 evolution is characterized by an increasing protein concentration along the TM region, indicating an enhanced tumor proteolytic activity, and by a steep profile ahead in the direction of tumor migration. Besides, we observe that the MMP1 external front is slightly shifted ahead of the tumor front region. Although MMP1 is produced along the protrusions, in fact, it is released in the extracellular space spreading in the areas around the tumor front, which perfectly fits with our results.

Finally, in Figure 6-*D* we show the situation after 15 hours of tumor evolution. Figures 6-*B*, 6-*C* and 6-*D* show that the invasion patterns of MMP1 and integrins are maintained over time. The experimental results in Figure 6-*DE* provide the distributions of GB membrane, active integrins, and protease markers at the same time, which co-localize in the TM region. The results also show a displacement of the MMP1 distribution with respect to active integrin and tumor membrane distributions. Our model predictions coincide with the experimental data in Figure 6-*DE*. The previous considerations about the relative positions of these three elements might be seen as an indicator of the direction of tumor migration.

As we developed in the Supplementary Text S2, the MMP1 activity modifies the porosity of the medium, facilitating spreading and, therefore, the transport and progression of the tumor mass. In particular, tumor cell advance velocity might not be constant, but linearly dependent on the porosity of the medium, that produces modifications of the tumor front, as shown in the Supplementary Figure S3. Furthermore, depending on the concentration of MMP1, even in case of constant tumor cell velocity, the front of the tumor may lose regularity and sometimes break into two different fronts. Then, if the tumor growth is not homogeneous and there is a certain heterogeneity in the growth of the invasion fronts, modifications and splittings of the frontal structure might arise, in agreement with experimental evidences (see Supplementary Figure S4).

## Discussion

The location of the GB advancement front is essential to predict its evolutionary dynamics and should help to design possible therapies.

In this work, we demonstrate that the tumor front is sharp and harbors a considerable biochemical activity. The protein ratio of integrins, proteases and focal adhesions defines the active front of GB. The evolution of these agents is also translated into sharp concentration fronts. At the same time, they produce biomechanical changes in the porosity and stiffness of the tissue, or in the infiltration capacity of the tumor mass located closer to the TM area.

A continuous feedback between experiments and modeling has found common ground in our analysis to advance knowledge on the dynamics of the GB front. Our results also confirm that *Drosophila* model represents a suitable platform to analyze the molecular and cellular mechanisms implicated in GB progression. Our mathematical model is based on the dynamical evolution and transport of the agents involved in the front progression. It has the ability to reproduce these biological features and, in particular, the evolution of GB invasion fronts, with a predictive value.

In conclusion, our results point out some of the main agents on the GB battlefront and also describe how mathematical models, incorporating this information, can predict its dynamics.

## Methods

### Experimental Procedures

#### Fly stocks

Flies were raised in standard fly food at 25 ° C. Fly stocks from the Bloomington stock Centre: UAS-myr-RFP (BL7119), repo-Gal4 (BL7415), tub-gal80ts (BL7019). Vienna Drosophila Resource Center (VDRC): UAS-*mys* RNAi (BL33642), UAS-*rhea* RNAi (BL28950). UAS-dEGFR*λ*, UAS-PI3K92E (dp110CAAX) (A gift from R. Read).

#### *Drosophila* glioblastoma model

To reproduce glioblastoma in *Drosophila* we took advantage of the binary expression system Gal4/UAS [50]. We over-expressed constitutively active forms of EGFR and PI3K under the control of UAS sequences. To restrict this expression to glial cells, we used the specific enhancer repo (repo-Gal4>UAS-EGFR*λ*, UAS-dp110).

#### Immunostaining

We dissected third-instar larval brains in phosphate-buffered saline (PBS), fixed in 4% formaldehyde for 25min, washed in PBS + 0.3% Triton X-100 (PBT), and blocked in PBT + 5% BSA. Antibodies: mouse anti-MMP1 (DSHB 1:50), mouse anti-FAK (Gift from Isabel Guerrero 1:50), rabbit anti-Talin (Gift from Isabel Guerrero 1:50), Secondary antibodies: anti-mouse Alexa 488, 568, 647, anti-rabbit Alexa 488, 568, 647 (Thermofisher, 1:500).

#### Imaging

*Drosophila* brain images were mounted in Vectashield mounting media with DAPI (Vector Laboratories) and analyzed by Confocal microscopy (LEICA TCS SP5) with a 63x oil immersion objective and 3x magnification. Images were processed using Image J 1.52t.

#### Quantifications

FAK, MMP1 and Talin signal was determined from images taken at the same confocal settings. Pixel intensity was measured using plot profile tool from Fiji 1.52t.

#### Survival

Animal survival is represented as the percentage of flies as compared to control siblings that reach adulthood upon GB induction (repo-Gal4>UAS-EGFR*λ*, UAS-dp110). n>100.

#### Numerical solution of the model

The whole system of partial differential equation (see (21) in the Supplementary Text S1), coupled with no-flux boundary conditions, was numerically solved with a self-developed code in Matlab (MathWorks Inc., Natick, MA). For the spatial discretization we considered the Galerkin method on a spatial grid of 500 points. In particular the flux-limited terms were discretized using an IMEX version of the Galerkin scheme. For the time discretization, an implicit Euler scheme was used for proteases and tumor equations, while a fourth order Runge-Kutta method for the other involved populations over a total of 15 · 10^6^ time points.

#### Analysis of the biological data

For the analysis of the experimental data concerning the distributions of MMPs, active and inactive integrins, we used a self-developed code in Matlab and the curve fitting toolbox, considering a weighted smoothing spline interpolation. In particular, for the analysis of MMP1 distribution we analyzed 26 measurements on 8 different images taken from as many animals; then, for the distribution of Talin and FAK, we analyzed 14 measurements on 6 different images taken from as many animals; finally, for the combined distribution of FAK and MMP1, we analyzed 23 measurements on 13 different images taken from as many animal.

## Acknowledgments

The authors thank Isabel Guerrero for the reagents, her valuable advice during the development of this work and for comments on the manuscript. The authors thank also Professors Luca Gerardo-Giorda, José L. López and Juan J. Nieto for their helpful discussions. This work has been partially supported by the MINECO-Feder (Spain) research grant number RTI2018-098850-B-I00 (JS), the Junta de Andalucía (Spain) Project PY18-RT-2422 & A-FQM-311-UGR18 (MC, SC, JS). This research is supported by the Basque Government through the BERC 2018- 2021 program and by the Spanish State Research Agency through BCAM Severo Ochoa excellence accreditation SEV-2017-0718. This project has received funding from the European Union’s Horizon 2020 research and innovation programme under the Marie Sk-lodowska-Curie grant agreement No. 713673. The project that gave rise to these results received the support of a fellowship from “la Caixa” Foundation (ID 100010434). The fellowship code is LCF/BQ/IN17/11620056. The authors contributed equally to the work. The authors declare no competing interests.

## Supplementary Text S1: Mathematical model

Our mathematical model is made up of five coupled differential equations for the description of the concentrations of MMPs (*P*), active (*A*) and inactive (*I*) integrins, as well as for the densities of glioma cells (*N*) and ECM (*E*). All these variables are functions of time (*t*) and space (*x*). In particular, since our objective is the study of the tumor front evolution, the differential equations for the different populations, and their numerical simulations, are formulated in a 1D case, i.e., (*t, x*) ∈ [0, *T*] × Ω, with *T* > 0 and Ω = [0, *b*_Ω_].

We develop the model considering that tumor evolution is characterized by a steep and well-defined front of invasion and an increased tumor activity in the front region. We refer to this region as TM region (*L*_*T M*_) and we characterize the heterogeneity of the tumor activity in the tumor domain using the functional ℱ(*N*). Defined the tumor *Sup*(*N*) = [0, *b*_*N*_] *⊆* Ω, the TM region is described as:

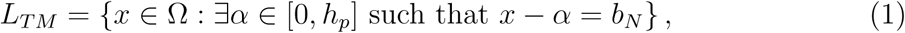

where *h*_*p*_ is the maximum length of a microtube. The functional for the tumor activity is given by:

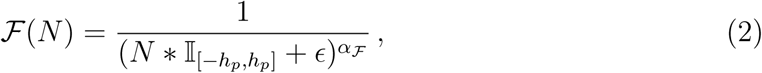

where *∗* indicates the convolution operator, 𝕀_[−*hp,hp*]_ represents the identity function on the interval of semi-amplitude *h*_*p*_, and *E* and *α*ℱ are parameters used for modulating the tumor activity. They are incorporated into the model on the base of the experimental results and these novelty features enable for a better fitting of the biological data.

### Glioma cell density (*N* (*t, x*))

The evolution of glioma cell density is described using the following equation:

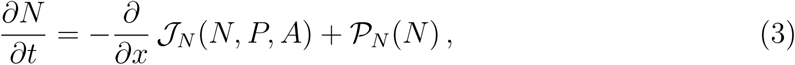

where 𝒥_*N*_ indicates the total flux of GB cells, and *𝒫*_*N*_ refers to the proliferation term. In particular, the flux 𝒥_*N*_ includes three parts: 𝒥_flux-sat_(*N*), 𝒥_chemo_(*N, P*) and 𝒥_hapto_(*N, A*), fluxes due to a flux-saturated mechanism, chemotaxis and haptotaxis, respectively. The operator describing the flux-saturated mechanism is defined as

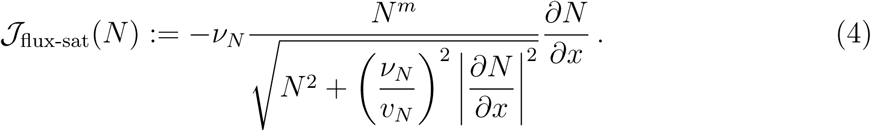

This type of operator was generally introduced with the so-called flux saturated equations in the form *∂*_*t*_*N* = −*∂*_*x*_ 𝒥_flux-sat_(*N*), where the exponent *m* is connected to the porosity of the medium [1]. These are equations in divergence form combining two non-linear diffusion mechanisms: the one of porous-media equations together with the one for the flux saturation mechanism, which provides a flow that is saturated as long as the size of the gradients is large enough. This type of equations is characterized by a finite speed of propagation bounded by *v*_*N*_, that limits the velocity of the propagation front and is incorporated in the model from experimental data. The velocity of the front depends also on the internal pressure, and it is exactly *v*_*N*_ for *m* = 1, while it is limited by *v*_*N*_ for *m* > 1 [2, 3]. *m* is also a parameter to fit from the experimental results. In this way, the solutions of the flux saturated equations preserve the characteristics of the initial data, in term of compactness of the support and possible jump discontinuities, allowing for the emergence of steep invasion profiles. Finally, the parameter *ν*_*N*_ represents the viscosity of the medium with respect to the movement of tumor cells.

Models with saturated flows appear in the pioneer works of Agueh, Brenier, Levermore, Mazon, Rosenau, Wilson, among others, in the literature on Astrophysics, wave propagation in a medium and in optimal mass transport as an alternative to linear diffusion (see the survey [3] for a historical overview). In fact, the latter fails to precisely define a propagation front, since it is characterized by the phenomenon known as infinite speed of propagation or instantaneous spreading. A model based on linear diffusion implies that GB cell would instantaneously reach and contaminate the whole brain tissue, to a greater or lesser extent, leading to the loss of the concept of tumor front and of the dynamics related to it. However, this is not commonly apparent. The capacity of tumor invasion and the possible response of the immune system makes necessary to focus our effort on controlling the battlefield. Therefore, the use of models based on saturated flux [2, 4] allows to locate and predict the dynamical evolution of the front.

The presence of MMPs that degrade the ECM, creating space for the tumor to migrate, determines a chemotactic flux given by:

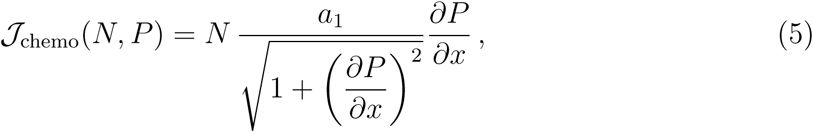

where *a*_1_ is the chemotactic sensitivity. This nonlinear form of the chemotactic flux reduces to the standard form under small gradients, while it saturates under large gradients.

The haptotactic flux 𝒥_hapto_, instead, is determined by the flux of active integrins that mediate the attachment process between GB and ECM. It is described as

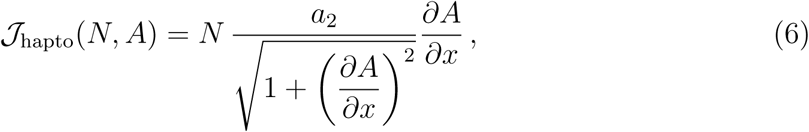

where *a*_2_ is the haptotactic sensitivity. Following the idea in [5], here the integrins produce an explicit effect on the direction of motion, describing the migration toward a gradient of recognized adhesion sites. By contrary with [5], we use a nonlinear form of the haptotactic gradient, analogous to the chemotactic one. Our novel approach is aimed at optimizing the influence on GB dynamics of the measurements of chemo- or hapto-forces along the trajectories, providing nonlinear terms in the corresponding Euler-Lagrange equations (5) and (6).

The proliferation term *𝒫*_*N*_ (*N*) simply describes a logistic growth of glioma cells at rate *a*_3_ and up to the maximum carrying capacity *K*_*N*_ :

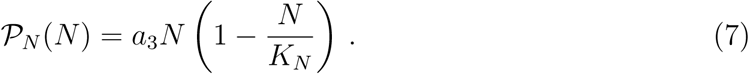

An improvement of this growth term should take into account the influence of morphogenic signaling pathways, such as Sonic Hedgehog (Shh) or Wnt (see [4, 6, 7, 8, 9, 10, 11] for further details). We will try to address this point in future works.

Combining the fluxes and the proliferation term leads to the following governing equation for glioma cells:

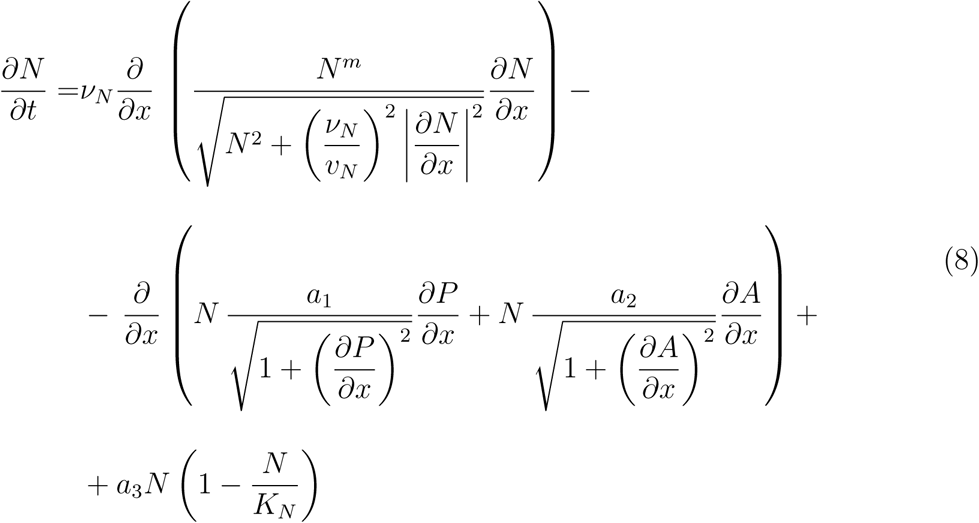

The combination of the flux-saturated mechanism and the chemo- and hapto-tactic fluxes might lead to different evolutions of the tumor profile, depending on the relative strength of the sensitivity parameters *a*_1_ and *a*_2_ and on the tumor velocity *v*_*N*_ and viscosity *ν*_*N*_. In the Supplementary Figure S4-A, we analyze how different values for the chemotactic sensitivity *a*_1_ might provide a separation of the tumor front. The variation of this sensitivity confirms what was experimentally observed (see, for instance, Supplementary Figure S4-D) about the fact that the tumor cell profile and the tumor proliferation are not always homogenous, and tumor cells might assume a strong proliferative phenotype in specific regions. Based on the variation of the parameters, the model predicts two possibilities. Cells close to the inner front start to proliferate more in order to fill and reduce the distance between the two developed tumor fronts, as shown in the Supplementary Figure S4-B. Or, instead, cell close to the outer front proliferate more in order to create an autonomous front, determining tumor profiles similar to the one shown in the Supplementary Figure S4-C. The heterogenous proliferation will be further investigated in future works, since this phenomenon has a major impact and relevance in an higher spatial dimension case.

### Metalloproteases concentration (*P* (*t, x*))

We suppose that MMPs are produced by tumor cells and released in the extracellular space to mediate the ECM degradation process. Their dynamics are described as follow:

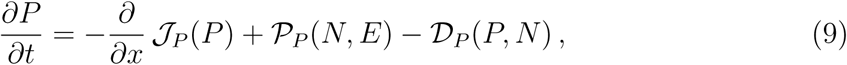

where 𝒥_*P*_ (*P*) is the MMP flux, *𝒫*_*P*_ (*N, E*) the production term and 𝒟_*P*_ (*P, N*) the degradation term.

The flux 𝒥_*P*_ (*P*) is described with the same flux-saturated mechanism used for the tumor cells, i.e.,

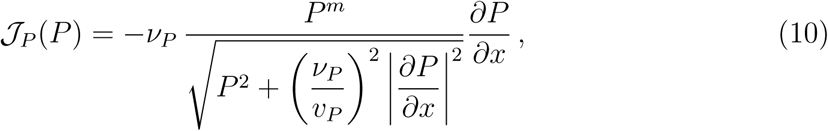

with *v*_*P*_ velocity of MMPs and *ν*_*P*_ viscosity related to MMP diffusion. The production term *𝒫*_*P*_ (*N, E*) is given by:

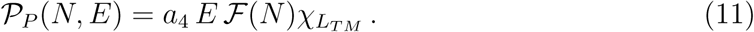

MMPs production is localized on the protrusion region and it is weighted using the tumor activity functional ℱ (*N*). MMPs are produced at rate *a*_4_, while 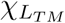 is the characteristic function of the TM region. Moreover, in the presence of ECM and once the tumor has moved forward, MMPs degrade at rate *a*_5_:

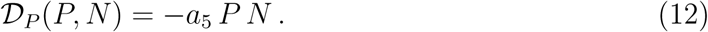

In particular, we assume that MMPs degradation preserve a basal level of proteins in the inner part of the tumor mass, estimated from the experiments. This is mainly related to the normal cell proteolytic activity in the main tumor mass, not directly aimed at sustaining the migration process.

The overall governing equation for MMPs is given by:

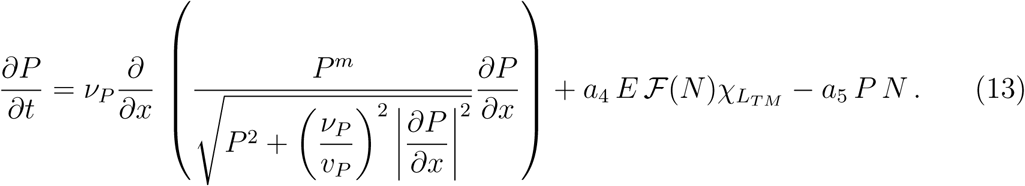

### Extracellular matrix (ECM) density (*E*(*t, x*))

For the extracellular matrix we model the degradation process due to the proteolytic activity as:

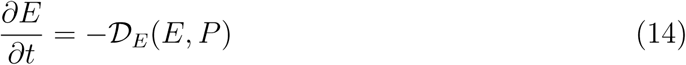

with degradation term 𝒟_*E*_(*E, P*) = *a*_6_ *E P*, and *a*_6_ degradation rate. Since, after the degradation process, some residual ECM material partially remains in the main tumor region, we include in the model a basal level of ECM inside the main tumor mass.

### Active and Inactive integrins concentration (*A*(*t, x*) **and** *I*(*t, x*))

We divide the integrin family into two subpopulations, active and inactive integrins, defining as active integrins those receptors which are actively bound to the ligands of the extracellular matrix. In particular, this subfamily determines the gradient responsible for the haptotactic movement of tumor cells. The corresponding equation is made up of three different terms: integrin activation, integrin inactivation and a flux term:

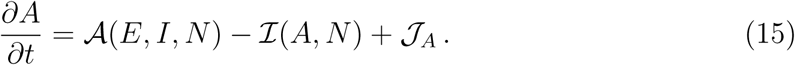

We model the binding between GB cells and ECM through binary interactions of inactive integrins and ECM at rate *a*_7_, weighting this process with the functional 𝒥_*N*_ :

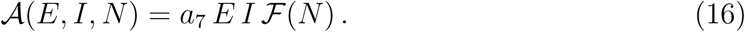

Instead, once the tumor has moved forward, inactivation occurs at rate *a*_8_ :

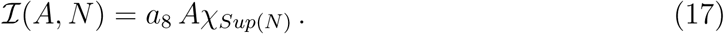

In particular, *χ*_*Sup*(*N*)_ is the characteristic function of the support of *N*. Active integrins are also subjected to a flux term describing the transport process due to the internal movement of GB cells. In fact, since integrin receptors are locate on the cell membrane, during the process of cell migration the receptors themselves are also transported:

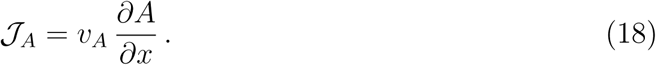

The transport velocity *v*_*A*_ is defined through a non-linear functional relationship that depends on the evolution of *N*, i.e., *v*_*A*_ = *v*_*A*_[*N*]. This functional describes the propagation rate of the support of N by means of its evolution equation.

The dynamic interactions between active and inactive integrins are demonstrated by the attachment and detachment terms. Furthermore, we consider that new inactive integrins are created at rate *a*_9_ and up to a maximum total level *K*_*I*_, due to an exocytosis process (*𝒫*_*I*_). Therefore, we define the following equation for inactive integrins dynamics:

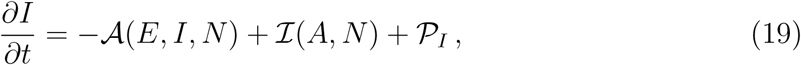

where

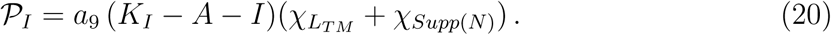

Our interest in the role of integrins and proteases as leading mechanisms of tumor progression arises from a serious of experiments that analyzed their activity and influence on tumor development. Integrin subunits *α*3, *α*6, and *α*7, for instance, are expressed in stem cell-like GB cells, they localize especially in invading cells, and mediate invasion [12], growth and survival [13], as well as proliferation [14] and GB formation [15]. They have emerged as an enrichment marker for brain tumor malignancy [13] and as a promising anti-glioblastoma target. Several data show the ablation of the invasive capacity in cells knocked down for integrin *α*7, in vitro and in vivo [13], the reduction of tumor formation for the functional knockdown of *α*6 signaling pathway [15], and the improvement of patients survival in combined therapy with standard chemoradiation [12]. Similar results come also from several clinical studies combining a broad spectrum of MMP inhibitors (for instance marimastat, that inhibits MMP-1,-2,-7,-9) with temozolomide, showing promising beneficial effects on the increment of progression-free survival (see [16] and reference there in).

The whole system of five coupled differential equations reads:

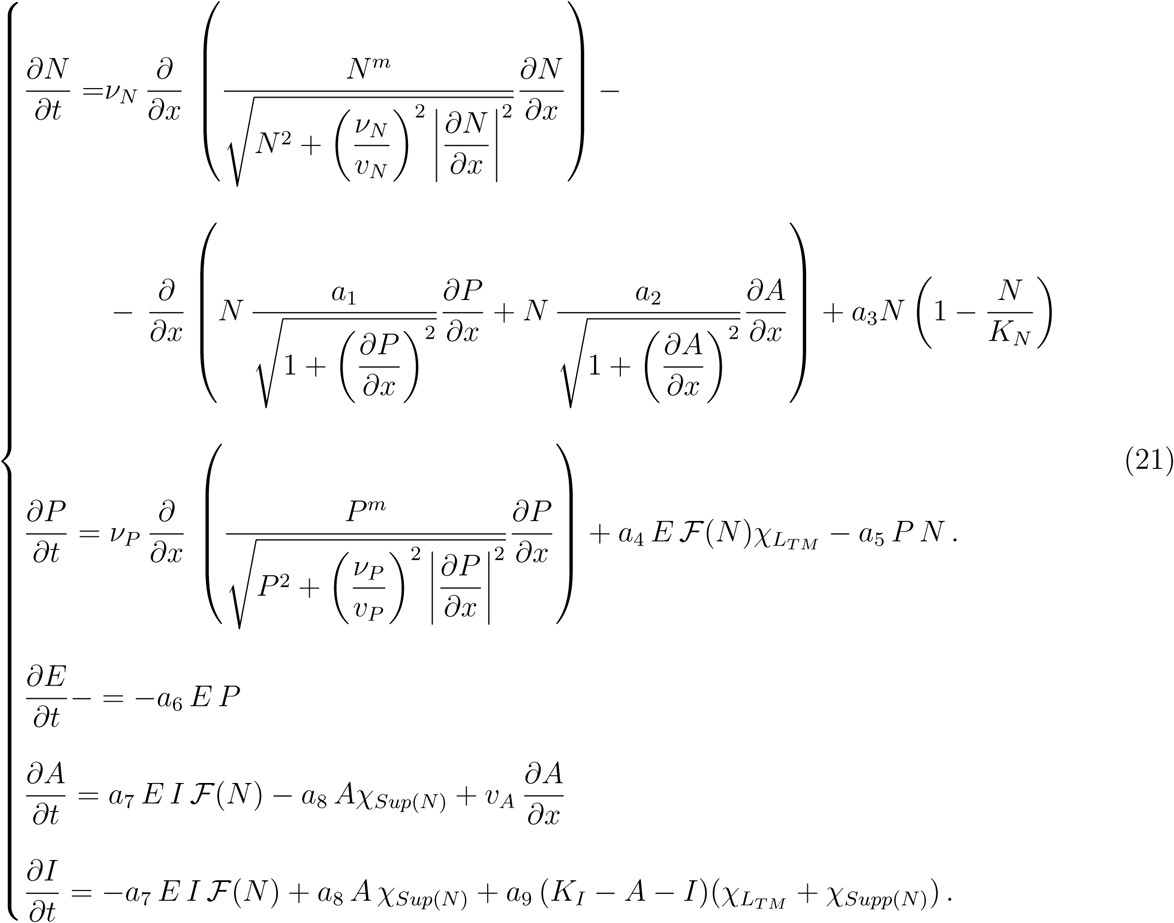

## Supplementary Text S2: Parameters estimation

We consider an undimensionalized version of system (21), scaling the five populations with respect to their carrying capacities (*K*_*N*_, *K*_*I*_) or to their typical concentrations/densities 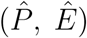. Table S1 reports the values of the model parameters and the reference values used for the undimensionalization procedure. For the numerical simulations presented in Figure 6 we use *m* = 1.

### Velocity and Viscosity parameters

The mean value reported in literature for glioma cell speed in human is 50 *µ*m · h^−1^ [25]. Since our biological experiments are done in a *Drosophila* glioblastoma model, we deduce the value for the tumor propagation velocity in *Drosophila* considering the Stokes law. It expresses the frictional force, also called drag force, exerted on spherical objects with very small Reynolds numbers in a viscous fluid. As a consequence, it deduces the relation between their velocity *v* and radius *R*:

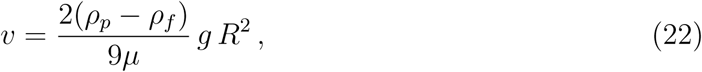

where *ρ*_*p*_ and *ρ*_*f*_ are the mass densities of the particles and the fluid, respectively, *µ* is the dynamic viscosity and *g* the gravitational field strength [26]. Assuming an average size of [12 − 14] *µ*m for human GB cells and 5 *µ*m for GB cells in *Drosophila* (measurements taken from our experiments), we deduce the range [6.4 − 8.7] *µ*m · h^−1^ for the parameter *v*_*N*_. Considering the strong relation between MMPs and GB cells, in terms of their respective locations and influences, and due to the lack of experimental data about protease propagation velocity in the brain, we assume *v*_*P*_ = *v*_*N*_.

For the tumor viscosity *ν*_*N*_, we refer to [4], where a similar description of the flux saturated mechanism is used for modeling the dynamics of the protein Shh. In [4], the authors consider Shh aggregates moving along a protrusion with speed *v*_*Shh*_ = 1.3·10^−3^ *µ*m · s^−1^ and kinematic viscosity *ν*_*Shh*_ = 5 · 10^−9^ cm^2^· s^−1^. Since in the description of the flux saturated operator a key role is played by the ratio 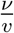, we use that 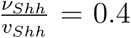mm, and we deduce *ν*_*N*_ ∈ [0.256 − 0.348] · 10^−2^ mm^2^· h^−1^.

To deduce the protease viscosity *ν*_*P*_, considering the similarity in size between MMP and Shh, we use the Einstein formula [27]. For a spherical particle of radius *R* moving with uniform velocity in a continous fluid of viscosity *µ*, the frictional coefficient is given by *f*_*τ*_ = 6 *π µ R*. Assuming that this applies also to spherical molecules, the kinematic viscosity is given by

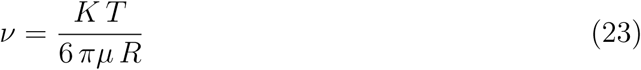

**Table S1:** Parameter estimation

| Parameter | Description | Value (units) | Source |
| --- | --- | --- | --- |
| $\nu_N$ | tumor viscosity | $[0.256 - 0.348] \cdot 10^{-2} \text{ (mm}^2 \cdot \text{h}^{-1})$<br>used value: 0.00348 | see description |
| $v_N$ | tumor velocity | $[0.64 - 0.87] \cdot 10^{-2} \text{ (mm} \cdot \text{h}^{-1})$<br>used value: 0.0087 | see description |
| $a_1$ | chemotactic sensitivity | 0.001 (mm<sup>2</sup> · h<sup>-1</sup>) | [17] |
| $a_2$ | haptotactic sensitivity | 0.0036 (mm<sup>2</sup> · h<sup>-1</sup>) | [17] |
| $a_3$ | tumor proliferation rate | 0.0345 (h<sup>-1</sup>) | [18] |
| $K_N$ | tumor carrying capacity | $\sim 10^6 \text{ (cells} \cdot \text{mm}^{-3})$ | see description |
| $\nu_P$ | MMPs viscosity | 0.035 (mm<sup>2</sup> · h<sup>-1</sup>) | see description |
| $v_P$ | MMPs velocity | $[0.64 - 0.87] \cdot 10^{-2} \text{ (mm} \cdot \text{h}^{-1})$<br>used value: 0.0087 | see description |
| $a_4$ | MMPs production rate | $[3.6 - 180] \text{ (h}^{-1})$<br>used value: 3.6 | [19] |
| $a_5$ | MMPs degradation rate | $[0.18 - 18] \text{ (h}^{-1})$<br>used value: 18 | [20] |
| $a_6$ | ECM degradation rate | $\sim 4.5 \text{ (h}^{-1})$ | [21, 22] |
| $a_7$ | integrin activation rate | $[1 - 3] \cdot 36 \text{ (h}^{-1})$<br>used value: 108 | [23] |
| $a_8$ | integrin inactivation rate | 36 (h<sup>-1</sup>) | [23] |
| $a_9$ | integrin exocytosis rate | $[0.36 - 36] \text{ (h}^{-1})$<br>used value: 0.72 | [24] |
| $K_I$ | integrins maximum capacity | $\sim 10^{10} \text{ (integrins} \cdot \text{mm}^{-3})$ | see description |
| $\hat{E}$ | ECM reference value | $10^{-3} \text{ (mg} \cdot \text{mm}^{-3})$ | [20] |
| $\hat{P}$ | protease reference value | $10^{-7} \text{ (mg} \cdot \text{mm}^{-3})$ | [20] |

with *K* Boltzmann constant and *T* temperature (in K). Considering a radius of *R*_*Shh*_ ∈ [10 − 100] nm for a vesicle containing Shh and moving along a cell protrusion, and using this information in equation (23), we get the following estimation

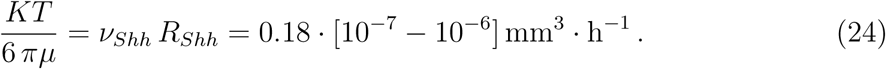

MMPs have a radius of 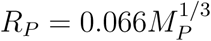 [28], with *R*_*P*_ expressed in nm and the mass *M*_*P*_ in Da. From [18], we know that *M*_*P*_ ∈ [72 − 92] kDa; therefore

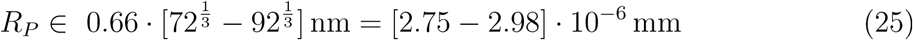

and

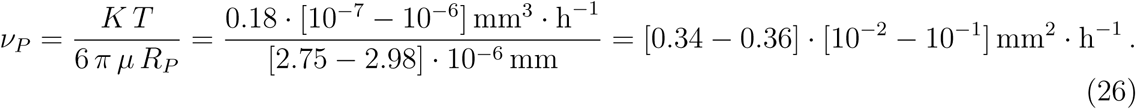

We set *ν*_*P*_ = 0.035 mm^2^ · h^−1^.

As the proteases degrade the ECM, the porosity of the medium increases. This process can be modeled in two different ways. By means of an equation for the degradation of the ECM (14) and considering its influence on the dynamics of the protease (13), and consequently on the spread of the tumor (8), which is the modeling approach we chose here. Or, an alternative way is to modify directly the porosity of the medium *E*, and to model its effect on the cell velocity. Several experiments [29, 30], in fact, have shown the relation between the cell velocity and the size of the pores of the ECM. Especially in absence of proteolytic activity, too dense ECM does not allow cells to move inside it, since the pores are too narrow with respect the cell capability of squeezing and passing through it. At the same time, too large pores do not allow for cell migration neither, since cell protrusions still need a certain amount of extracellular matrix around them in order to attach to it. Following the results of [30] (see, for instance, Figure 2.b in there), we consider a variability range for *ϵ* ∈ [0.5 − 0.75], and we define a law of variability for *v*_*N*_, assuming an optimal values of the tumor cell velocity for *ϵ* ∈ 0.67. Using an evolutionary law for *ϵ* analogous to the one proposed in [31]

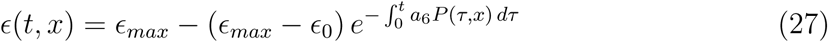

with *ϵ*_0_ = 0.54 initial porosity value, and *ϵ*_*max*_ = 0.75, we test our model for a scenario of non-constant *v*_*N*_. Results are shown in the Supplementary Figure S3.

### Carrying capacities

The carrying capacity of tumor cells is estimated considering the mean diameter of a GB cell in *Drosophila* (5 *µ m*). This leads to an order of magnitude for the carrying capacity of *K*_*N*_ *∼* 10^6^ cells · mm^−3^.

Instead, for the integrin capacity, considering that there are *∼* 10^5^ integrin receptors per cell [32], we estimate a maximum of *K*_*I*_ *∼* 10^10^ integrins · mm^−3^.

## Supplementary Figures and Comments

To validate the functional contribution of integrins to GB progression we use specific RNAi constructs to knockdown *myospheroid (mys)* or *rhea*, two key players for integrin function. *mys* encodes a *β* subunit of the integrin dimer that acts as adhesion/signaling protein, regulating cellular adhesion and migration. Besides, *rhea* encodes *Drosophila* Talin, a large adaptor protein essential for adhesive functions of integrins. Confocal images of *Drosophila* brains with GB show large brains with expanded GB cell membrane (red Figure S1-*A*). Upon *mys* or *rhea* knockdown, GB expansion is impaired (Figure S1-*B* and *C*) and the lethality caused by GB is partially rescued (Figure S1-*D*). These results suggest that GB cells require intact integrin function to progress, expand and cause premature death.

We quantified Talin and FAK immunostaining signals in confocal microscopy GB images to validate the inverse relation of Talin and FAK in inner GB mass and at GB front. The results show that the inner region of the GB mass (Figure S2-*A*_1_) has higher Talin protein levels (Figure S2-*A*_2_) and lower FAK signal (Figure S2-*A*_3_), as represented in Figure S2-*A*_4_. In line with our previous results, the relative concentration of Talin and FAK is inverted in the front region of GB samples (Figure S2-*B*_1_). The results, in fact, indicate that Talin concentration drops significantly (Figure S2-*B*_2_ and *B*_4_) and correlates with an increase of FAK signal at GB front (Figure S2-*B*_3_ and *B*_4_). These observations suggest that Talin and FAK maintain an inverse correlation, and they are indicators of the migratory status of GB cells.

**Figure S1:**
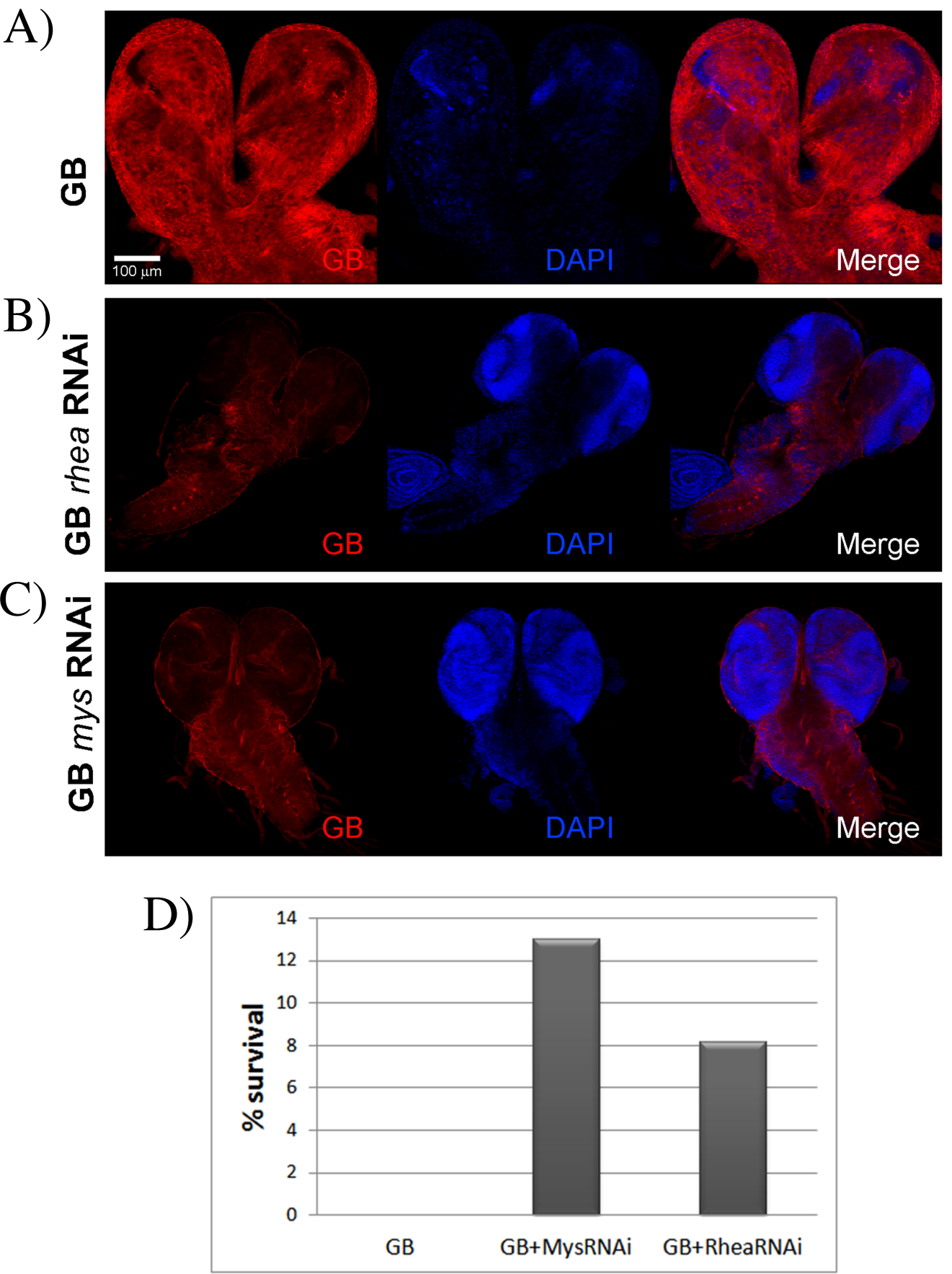
Functional integrins are required for GB progression. Low magnification confocal images of *Drosophila* GB larvae brain in A) and *rhea* knockdown (UAS-*rhea* RNAi) in B), or *mys* knockdown (UAS-*mys* RNAi) in C). GB cell membrane is marked with myristoylated-RFP (red) and nuclei are marked with DAPI (blue). The images show that *rhea* or *mys* knockdown prevents the expansion of GB (red) and rescues the lethality (percentage of survival in D).

**Figure S2:**
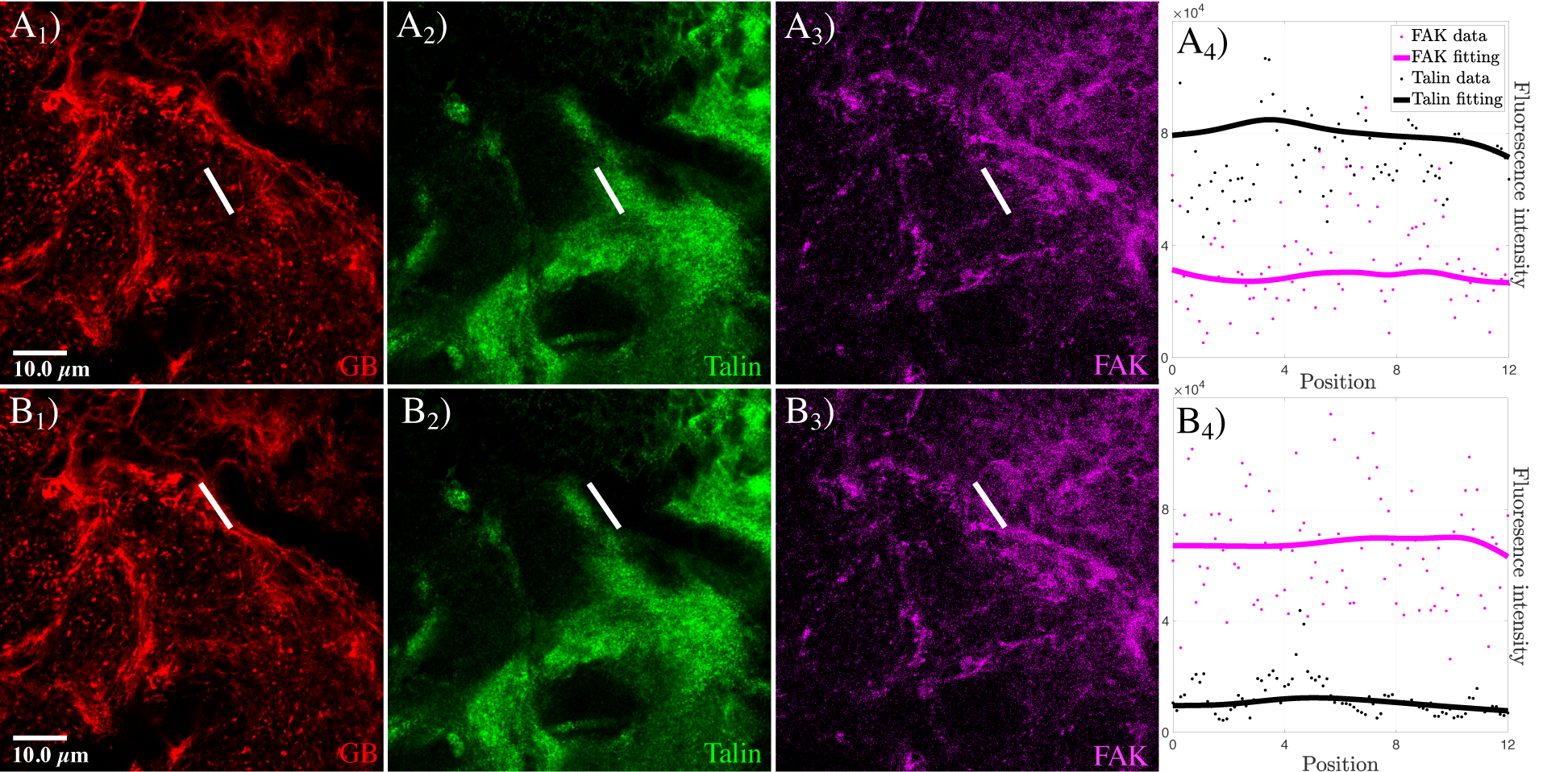
Talin and FAK dynamics: comparison between the inner GB mass and the GB front. Fluorescent confocal images of *Drosophila* 3rd instar larvae brain with GB marked in red (*A*_1_ and *B*_1_), and stained with anti-Talin in green and anti-FAK in magenta (*A*_2_, *A*_3_, *B*_2_ and *B*_3_). In *A*_4_ and *B*_4_, quantification of the fluorescent signals and graphical representation of the fluorescent intensity for GB, Talin and FAK signals along the white lines shown in *A*_1_-*A*_3_, *B*_1_-*B*_3_. Dots represent the data and lines represent the fitting.

**Figure S3:**
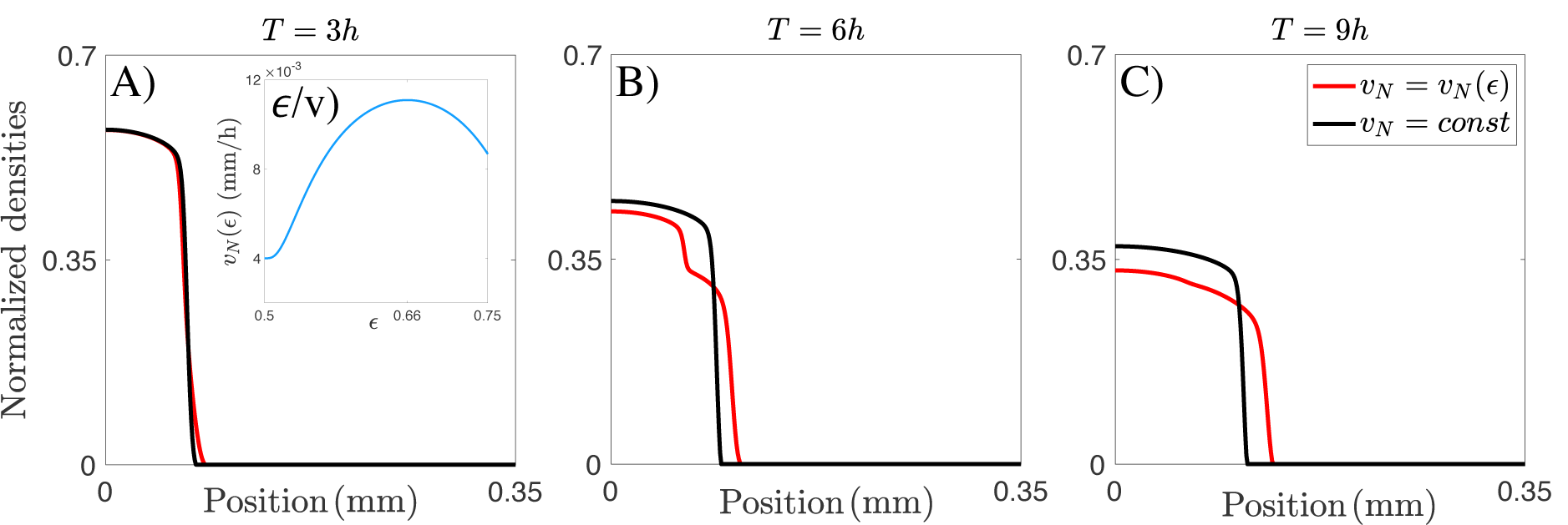
Effects of porosity changes on tumor profile. *A*), *B*), and *C*) show the comparison between the tumor density profile in the case of flux saturated model with constant velocity *v*_*N*_ (in black) and with *v*_*N*_ = *v*_*N*_ (*ϵ*) (in red) at the time steps referring to 3, 6 and 9 hours of tumor evolution, respectively. For this comparison we consider a tumor cell equation given by 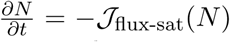. In *ϵ/v*), the profile of *v*_*N*_ = *v*_*N*_ (*ϵ*) is shown. In particular, accordingly with [30], the minimum value for the velocity relates to a porosity of 50%, while the maximum occurs around the value of 66%. The red curve shows how, as the velocity changes due to the ECM degradation process, that increases the medium porosity, cells closer to the front start moving faster than inner cells. This determines an heterogenous modification of the invasion front, that slightly exceeds the homogenous front of the constant velocity case. Eventually the entire main tumor mass feels the changes in the velocity and a unique front is recovered. If there is heterogeneity in the growth of the front, the profile might not unify and a new front might emerge from this disturbance, as in the Supplementary Figure S4-*C*).

**Figure S4:**
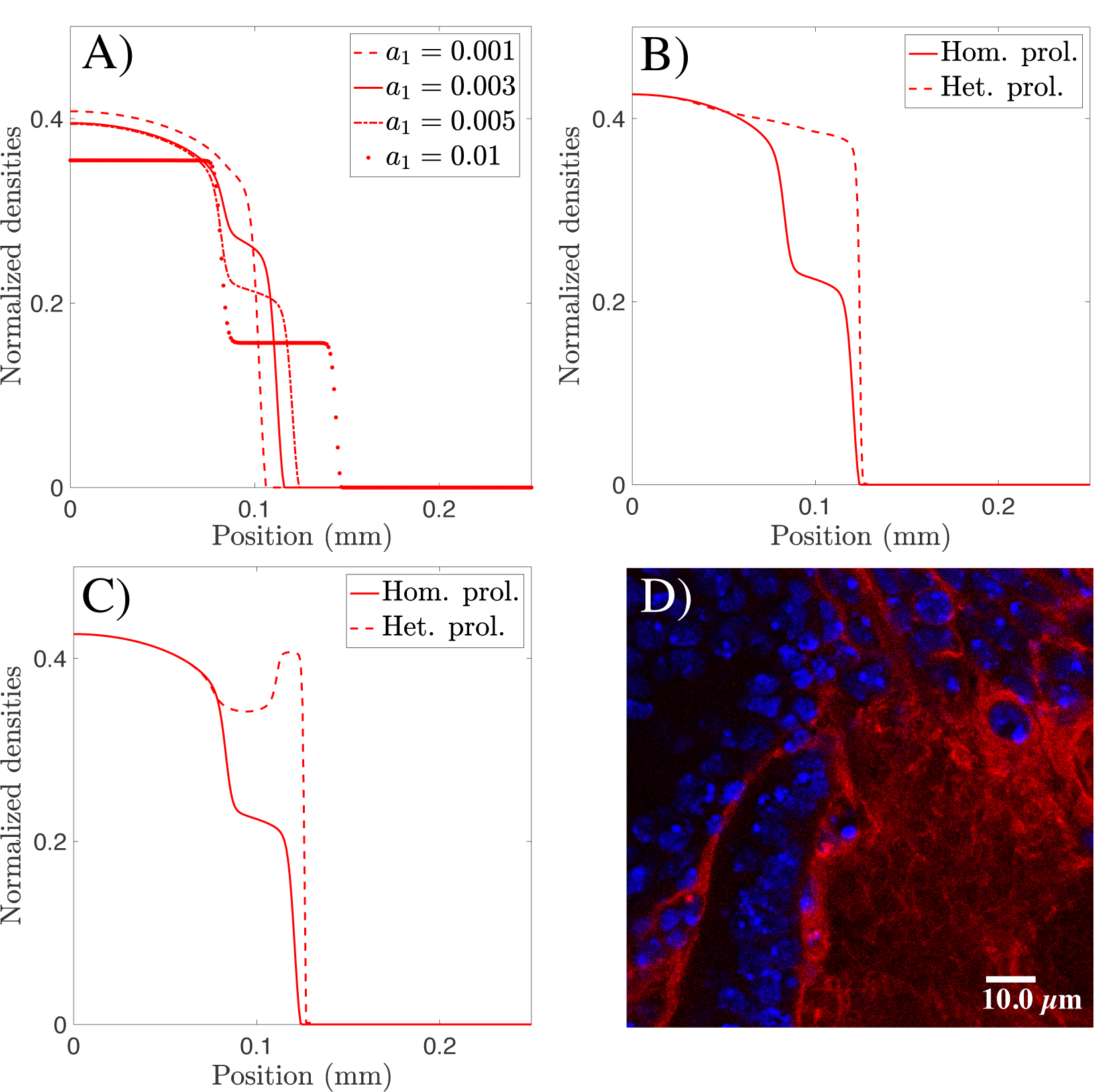
Effects of chemotaxis strength and heterogenous proliferation on tumor profile. Figure *A*), *B*) and *C*) show the evolution of the tumor profile after 5 hours in different cases. In *A*), the effect of changes in the chemotactic sensitivity *a*_1_ is presented. Since the proteolytic activity and, consequently, the concentration of MMP1 is enhanced in the front area, the stronger the parameter *a*_1_ is, the more evident the localization of tactic effect is. Tumor cells closer to the front acquire an increased overall velocity (due to both ℱ_flux-sat_ and ℱ_chemo_ fluxes) that leads to heterogenous fronts and, eventually, it might leads to a break of the tumor in two separated masses, as it can be observed in the experimental Figure *D*), where cell nuclei are marked in blue with DAPI, while GB cell membrane is marked with mystoylated-RFP in red. In particular, in *D*) higher intensity of the GB membrane marker indicates areas of tumor invasion. *B*) and *C*) show the comparison of our model with classical homogenous proliferation with the two possible models for heterogenous proliferation, in the case of *a*_1_ = 0.005.

